# Loss of connexin 43 in cartilage causes mitochondrial dysfunction and accelerates post-traumatic osteoarthritis progression

**DOI:** 10.64898/2026.06.03.728771

**Authors:** Kristyna Hargitaiova, Rebecca M. Irwin, Khizar Hayat, James Pham, Claire Ma, Alexandria M. Davis, Miguel Otero, Michelle L. Delco

**Affiliations:** Cornell University, Department of Clinical Sciences, Ithaca, NY; City University of Hong Kong, Hong Kong; Hospital for Special Surgery and Weill Cornell Medical College, New York City, NY

**Keywords:** Connexin 43, post-traumatic osteoarthritis, osteochondral communication, crosstalk, mitochondrial bioenergetics

## Abstract

Osteoarthritis (OA) is a major cause of chronic pain and disability worldwide, characterized by progressive degeneration of cartilage and subchondral bone. Post-traumatic OA (PTOA) develops in as many as 25–50% of individuals following major joint injury, making it a leading cause of OA in younger and otherwise healthy populations.^1,2^ Connexin 43 (Cx43), a gap junction protein involved in intercellular communication and cellular stress responses, has been linked to OA; however, its role in the progression of PTOA remains unclear. Here, we examined how cartilage-specific loss of Cx43 influences PTOA and chondrocyte metabolic function. Using a murine model of conditional Cx43 deletion in cartilage, we demonstrate that male knockout mice exhibited severe cartilage surface damage and matrix loss, whereas female knockout mice showed cartilage thinning accompanied by reduced chondrocyte hypertrophy, decreased subchondral bone density, and increased osteophyte formation. Thus, loss of Cx43 disrupts cartilage integrity and osteochondral remodeling in a sex-specific manner, predisposing joints to maladaptive bone changes and cartilage degeneration. Complementary mechanistic studies in human articular chondrocytes revealed that Cx43 deficiency impairs mitochondrial respiration, reduces spare respiratory capacity, and lowers ATP production, consistent with compromised cellular bioenergetics. Together, these findings identify Cx43 as an important coordinator of metabolic and structural responses to joint injury. These results position Cx43 as a context-dependent regulator of joint homeostasis and suggest that maintenance of Cx43 expression may support cartilage resilience following injury.

## Introduction

Osteoarthritis (OA) is one of the most prevalent and disabling musculoskeletal disorders worldwide, affecting over 300 million people and ranking among the leading causes of years lived with disability globally.^3^ Post-traumatic OA (PTOA), a common consequence of joint injury, disproportionately affects younger, active individuals^4^ and accounts for ~12% of all symptomatic OA.^2^ Despite decades of research and substantial investment in OA therapeutic development, no disease-modifying osteoarthritis drugs (DMOADs) have been approved to date and treatment strategies remain primarily limited to symptom management and joint replacement.^5,6^ This therapeutic gap reflects, in part, an incomplete understanding of the molecular drivers that govern early disease initiation and progression following joint injury.

Although articular cartilage is avascular and relies primarily on glycolysis rather than mitochondrial oxidative phosphorylation for energy production during homeostasis, mounting evidence implicates mitochondrial dysfunction as an early and central driver of OA pathogenesis.^7–9^ Recent work has highlighted the role of chondrocyte mitochondria as mechanotransducers during cartilage injury.^10–12^ Mechanical overload of the articular surface caused rapid, strain-dependent mitochondrial depolarization, impaired oxidative phosphorylation, excess reactive oxygen species (ROS) production, apoptosis, and catabolic gene expression prior to detectable cartilage loss, implicating mitochondrial dysfunction in the initiation of PTOA.^7,13^ Furthermore, dysregulated mitochondrial quality control - manifesting as disrupted fission/fusion and impaired mitophagy - was found to exacerbate inflammation and catabolic signaling, thereby accelerating cartilage degeneration.^8^

Articular chondrocytes are not isolated within individual lacunae but are interconnected by long filapodial extensions that traverse the dense extracellular matrix to form three-dimensional cellular networks.^14^ At cell-cell junctions, chondrocytes are linked via gap junction channels, comprised mainly of connexin 43 (Cx43), encoded by the *GJA1* gene. In many tissues, gap junctions are essential in maintaining homeostasis, allowing direct cell-cell signaling and synchronized regulation of nutrients and metabolites, although their role in articular cartilage is not completely understood.^15^

Cx43 acts as a mechanoresponsive protein in musculoskeletal tissues, modulating both chondrocyte and osteocyte intracellular signaling in response to mechanical loading.^16–18^ Further, increased Cx43 expression is associated with elevated levels of p53, p16 INK4a, SASP factors, COX-2, IL-1β, and cartilage matrix-degrading enzymes^19^, while down-regulation of Cx43 reversed pro-senescent and pro-inflammatory gene expression in OA cartilage.^20^ While these findings establish a connection between Cx43 and disease progression in OA cartilage, the recently discovered non-canonical functions of *GJA1* in supporting mitochondrial activity and protective stress responses are relatively unexplored in OA pathology.

In non-musculoskeletal cell types, the alternatively translated, 20kD N-terminal truncated isoform of Cx43 (GJA1-20k) protected against cardiac ischemia-reperfusion injury by promoting mitochondrial biogenesis and quality control.^21,22^ We recently demonstrated that Cx43 mediates intracellular mitochondrial transfer from mesenchymal stromal cells (MSCs) to injured articular chondrocytes, improving cellular bioenergetics and viability, and identified a novel role for GJA1-20k in MSC-chondrocyte mitochondrial transfer under oxidative stress conditions.^23^ These findings suggest a stress-responsive mechanism by which Cx43 may support recovery following injury, specifically through mitochondrial pathways. However, the effects of Cx43 on chondrocyte mitochondrial function and its role in cartilage during PTOA development have not been investigated.

Therefore, we developed a cartilage-specific Cx43 knockout (cKO) mouse model and evaluated PTOA progression after surgical destabilization of the medial meniscus (DMM). We report that Cx43 cKO exacerbates PTOA progression in a sex-specific manner by altering osteochondral remodeling, resulting in maladaptive bone changes and accelerated cartilage degeneration. Mechanistically, we examined mitochondrial activity and energy metabolism in human Cx43-knockout chondrocytes. Cx43 loss impaired mitochondrial respiration, reduced spare respiratory capacity, and decreased ATP production, indicating compromised chondrocyte bioenergetics. Together, these findings indicate that Cx43 coordinates metabolic and structural responses to joint injury, and highlight Cx43 as a context-dependent regulator of joint homeostasis, suggesting that Cx43 may support cartilage resilience following joint injury.

## Results

### Gja1 is efficiently deleted in articular cartilage of Cx43^cKO^ mice

To validate efficient and cartilage-specific deletion of Cx43 following tamoxifen (TMX) induction, we assessed *Gja1* levels in joint tissues from Cx43^cKO^ mice and wild-type (Cx43^WT^) counterparts. Immunohistochemistry (IHC) performed on whole murine knee joint sections confirmed loss of Cx43 protein in articular cartilage and meniscus of Cx43^cKO^ mice at 7 days post-TMX administration. In articular cartilage, Cx43 protein staining was reduced in Cx43^cKO^ compared to Cx43^WT^ mice (two-tailed Mann-Whitney test, median Δ = −12.3 % of surface area, U = 0, p < 0.0001, n = 16 cKO, n = 9 WT, Figure 1C, E). Similar results were observed in meniscus tissue, where Cx43^cKO^ knees had lower Cx43 staining compared to Cx43^WT^ (median Δ = −15.6 % of surface area, U = 0, n = 8cKO, n = 5 WT, p = 0.002, Figure 1C, E).

**Figure 1.**
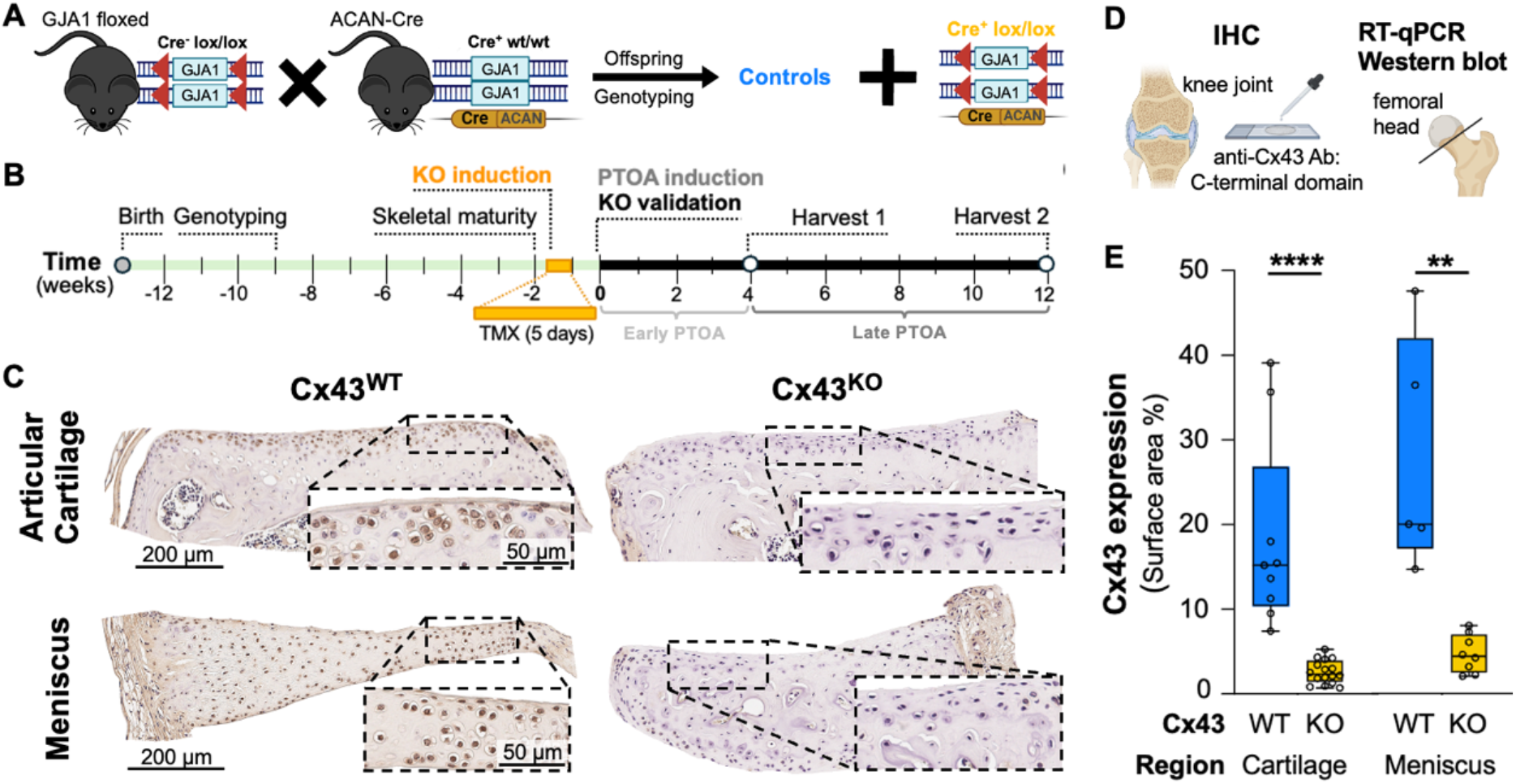
Study Design, generation and validation of tissue-specific Cx43 knockout in cartilage. A) Conditional Gja1 knockout (Cx43^KO^) mice were generated by crossing Gja1flox/flox mice with Acan-CreERT2 mice. B) Study Timeline; Tamoxifen (TMX, 75 mg/kg/day × 5 days) was administered at skeletal maturity (3 months of age) to induce cartilage-specific Cx43 deletion. C-E) KO validation was performed 7 days post-TMX injection by C) immunohistochemical staining for Cx43 in the articular cartilage of the medial tibial plateau (top row) and meniscus (bottom row) of Cx43^WT^ (left) and Cx43^KO^ (right); Insets highlight Cx43 staining (brown) within chondrocytes and fibrochondrocytes, respectively. E) Quantification of Cx43 staining in articular cartilage and meniscus. Cx43^KO^ mice exhibited significantly reduced Cx43 protein in cartilage and meniscus. Each dot represents one biological replicate (individual animal, n = 5-16 per group); horizontal bars indicate median, **p < 0.01, ****p < 0.0001.

Quantitative PCR analysis further supported efficient cartilage-specific Cx43 deletion (Figure S1). In femoral head cartilage harvested at 12-weeks post-injury (6-months of age), *Gja1* expression was significantly reduced in Cx43^cKO^ compared to Cx43^WT^ controls (one-tailed unpaired t-test, t(6) = 2.04, p = 0.04; mean difference = −0.38 ± 0.19; n = 4 per genotype), but not in control tissue types (heart and kidney). Expression of the chondrogenic marker aggrecan (*Acan)* was not different between genotypes in either articular or xiphoid cartilage (Figure S1). For additional knockout validation details, see *Supplemental Materials*.

### Cartilage-specific Cx43 knockout exacerbates PTOA in a time- and sex-dependent manner

To determine how cartilage-specific loss of Cx43 influences PTOA progression, we induced OA using the DMM model in male and female Cx43^cKO^ and Cx43^WT^ mice and evaluated joints at 4- and 12-weeks following DMM, corresponding to early and established OA, respectively (Figure 2A-C).^24–26^ Histological evaluation using the OARSI scoring system^27^ confirmed the expected DMM-induced PTOA-like changes in control (Cx43^WT^) mice in both sexes (Figure 2E), consistent with previous reports.^28–30^ In males, OARSI total joint scores did not differ between Cx43^WT^ and Cx43^cKO^ at 4 weeks post-DMM (main effect of genotype: F(1,17) = 0.27, p = 0.61; Figure 2F). However, at 12 weeks post-injury, male Cx43^cKO^ mice exhibited higher OARSI scores than Cx43^WT^ controls, with a main effect of genotype observed across both sham and DMM joints (mean difference (Δ) = −4.2, p = 0.007 both, genotype effect: F(1,17) = 14.48, p = 0.0014; Figure 2F), indicating more severe disease in the late PTOA phase. In females, divergence between genotypes occurred earlier; at 4 weeks post-DMM, Cx43^cKO^ females already exhibited higher OARSI scores than Cx43^WT^ in both sham- and DMM-limbs (genotype effect: F(1,18) = 9.2, p = 0.007; Δ = −3.34; 95% CI: −5.66 to −1.03 for both, Figure 2G), a difference that persisted at 12 weeks in the DMM limb (genotype effect: F(1,18) = 6.5, p = 0.02, Δ = −3.6; −6.6 to −0.6p = 0.02, Figure 2G).

**Figure 2.**
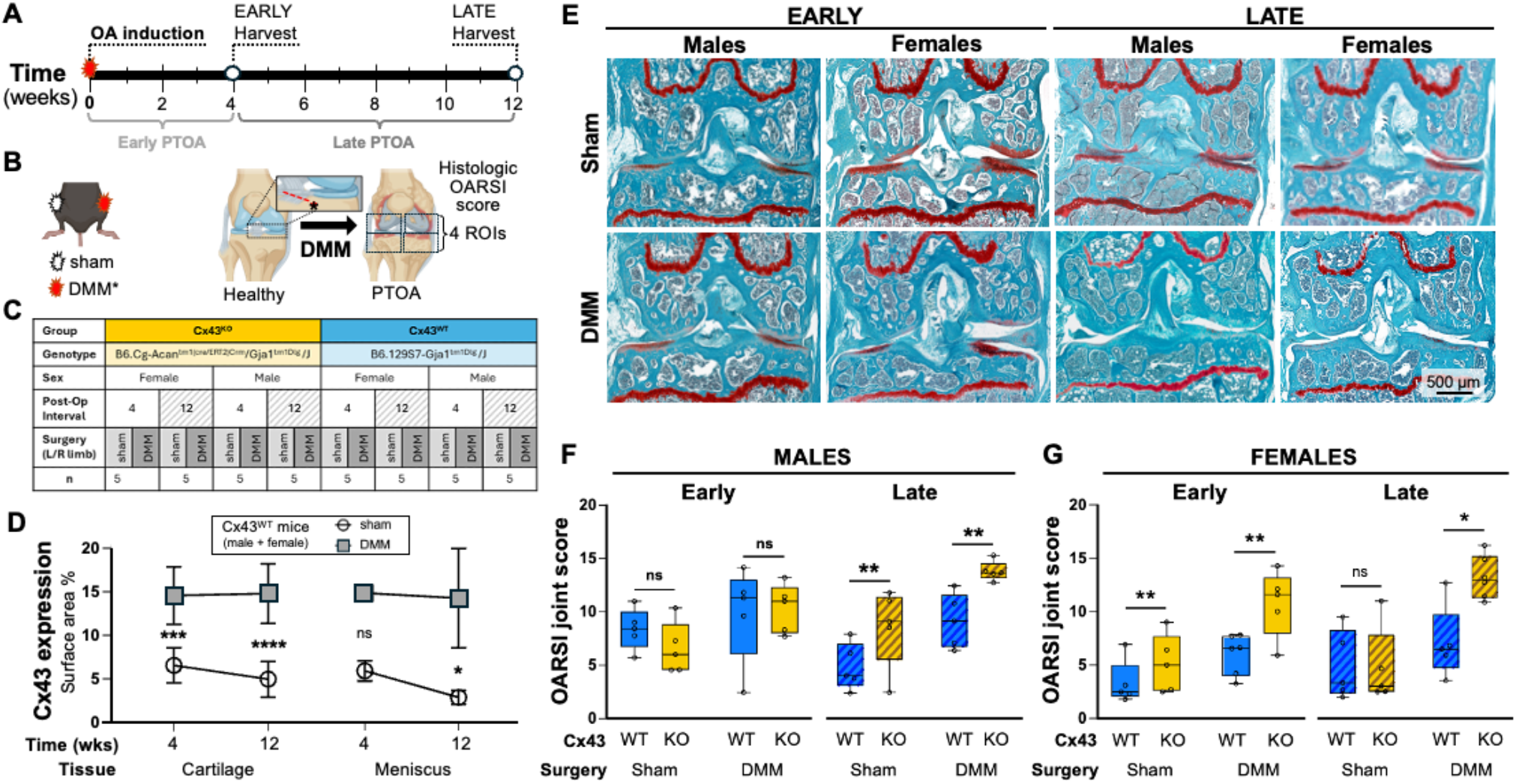
Histological assessment of joint degeneration in DMM and sham limbs at 4 and 12 weeks post-injury. A) Experimental Timeline with histology endpoints at early (4 weeks) and late (12 weeks) post-traumatic osteoarthritis (PTOA) stages. B) Methods for PTOA induction, with schematics representing the destabilization of the medial meniscus (DMM) surgical technique and contralateral sham (arthrotomy only) procedure (left, center), and histological regions of interest (ROI) for OARSI joint scoring (right). C) Summary table of experimental groups. D) Cx43 expression in articular cartilage was higher in DMM limbs compared to sham at both 4 weeks (p = 0.0007) and 12 weeks (p < 0.0001), with no change over time. In the medial meniscus, Cx43 expression did not differ between sham and DMM limbs at 4 weeks but was higher in DMM limbs at 12 weeks (p = 0.01). E) Representative whole-joint coronal histologic sections (saf-O/Fast green staining) from male and female Cx43^WT^ mice depicting progressive cartilage degeneration in DMM and sham limbs over time (early timepoint; left, late timepoint; right). F) In males, no differences in PTOA severity were observed between genotypes in sham or DMM limbs at 4 weeks (early); however, Cx43^KO^ DMM limbs had worse PTOA (higher OARSI scores) at 12 weeks (p = 0.02). G) Females displayed worse PTOA in DMM limbs at both early (p = 0.005) and late (p = 0.004) timepoints. Data (n = 5) was analyzed using two-way ANOVA with surgery and genotype as fixed factors, and is displayed as mean ± SD. *p<0.05, **p<0.01, ***p<0.001, ****p<0.0001.

Cx43 IHC analyses revealed a significant increase in Cx43-positive area in WT articular cartilage at 4- (Δ = −8 % surface area, 95% CI: −12.1 to −4.0; p = 0.0007) and 12-weeks (Δ = −9.8% surface area, 95% CI: −13.7 to −6.0; p < 0.0001, Figure 2D) following DMM vs. sham-operated limbs. A similar trend was seen in the medial meniscus at both the early (Δ = −8.9% surface area, 95% CI: −18.5 to 0.6; p = 0.06) and late timepoints (Δ = −11.3% surface area, 95% CI: −20.4 to −2.4; p = 0.02) post-injury (Figure 2D). In both cartilage and medial meniscus, the effect of DMM surgery was significant (cartilage: F(1,15) = 47.23, p < 0.0001, meniscus: F(1/15) = 10.84, p = 0.005), indicating injury increased Cx43 protein in joint tissues. Data was analyzed by mixed-effects modeling and are reported as mean differences and 95% CI.

### Cx43 deficiency drives sex-divergent cartilage responses during early PTOA

Because standard OARSI scoring^24^ does not distinguish between specific structural and cellular features of osteoarthritis, we also performed a feature-level histological analysis using a composite scoring rubric, adapted from published murine OA histopathology studies^24,26–29,31,32^ to separately assess cartilage surface integrity, thickness, hypertrophic chondrocyte (HC) abundance, bone lysis and sclerosis and periarticular osteophytes (Table 1).^29,30^

**Table 1.** Histological scoring rubrics for assessing osteoarthritis features.

| Modified OARSI Scoring Rubric |  |  |  |  |  |  |  |  |
| --- | --- | --- | --- | --- | --- | --- | --- | --- |
| Score | 0 | 0.5 | 1 | 2 | 3 | 4 | 5 | 6 |
| OA severity | Normal | Loss of Safranin-O without structural changes | Small fibrillations without loss of cartilage | Vertical clefts down to middle layer (immediately below superficial) | Vertical clefts down to calcified cartilage extending to < 25% | Vertical clefts down to calcified cartilage extending to 25-50% | Vertical clefts down to the calcified cartilage extending to 50-75% | Vertical clefts down to calcified cartilage extending to >75% |
| Modified Composite Scoring Rubric |  |  |  |  |  |  |  |  |
| Score / Parameter | 0 | 1 | 2 | 3 | 4 |  |  |  |
| GAG loss | No loss | < 25% | 25-50% | 50-75% | < 75% |  |  |  |
| Hypertrophic chondrocytes | None | Few (rare) | Mild (occasional) | Moderate (some) | Severe (many) |  |  |  |
| Surface architecture | Normal | Mild Irregularity (small fibrillations without cartilage loss) | Moderate irregularity (larger fibrillations without cartilage loss) | Severe irregularity (large fibrillations with focal cartilage loss) | Severe/worst irregularity (large fibrillations, large areas of cartilage loss) |  |  |  |
| Cartilage thickness | Normal | Mild thinning | Moderate thinning | Severe thinning | Single layer of cartilage tissue |  |  |  |
| SCB lysis | None | Minimal | Mild | Moderate | Severe |  |  |  |
| SCB sclerosis | None | Minimal | Mild | Moderate | Severe |  |  |  |
| Osteophytes | None | Minimal | Mild | Moderate | Large |  |  |  |
| Growth plate stain | Normal stain | < 25% GAG loss | 25-50% GAG loss | 50-75% GAG loss | > 75% GAG loss |  |  |  |
| Subjective OA score | Normal to minimal changes | Mild OA | Moderate OA | Advanced OA | Severe OA |  |  |  |

At 4 weeks post-DMM, sex-dependent responses were evident (Figure 3). At the structural level, total (whole-joint) cartilage Surface Damage scores were significantly affected by genotype x sex interaction (F(1,16) = 13.16, p = 0.002). Cx43^cKO^ males displayed worse surface damage compared to WT males (Δ = 4.9, 95% CI: −7 to −2.9, p < 0.0001) and KO females (Δ: 3.1, 95% CI: 1.02 to 5.18 p = 0.006), whereas no genotype-dependent difference was observed in females (Figure 3B). Males also had worse Cartilage Thinning scores compared to Cx43^WT^ males (Δ: −4.1, 95% CI: −6.95 to −1.32, p = 0.007), while no differences in cartilage thickness were detected between KO and WT females (p = 0.7; Figure 3C) at the 4-week timepoint.

**Figure 3.**
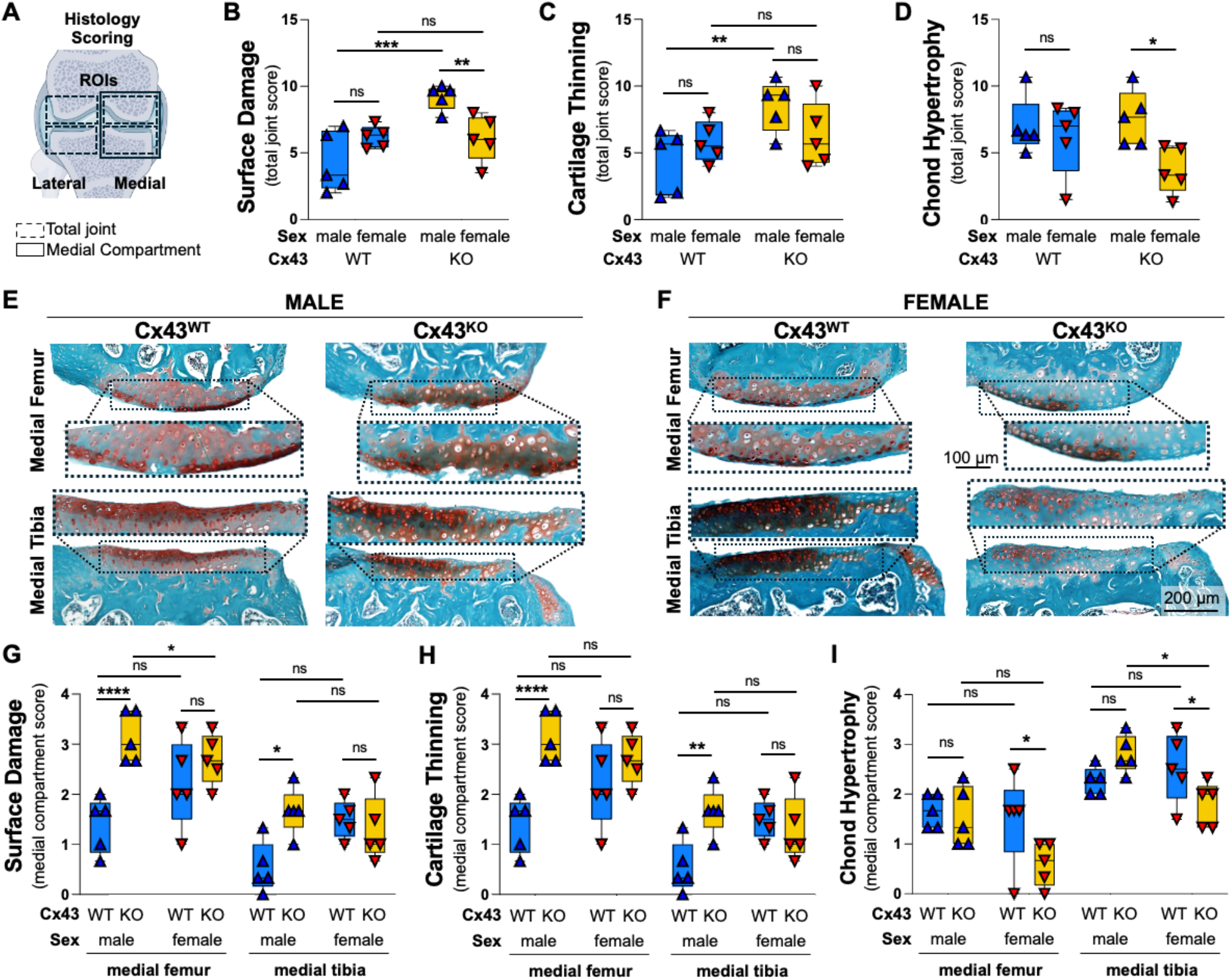
Cx43 cartilage-specific knockout drives sex-divergent cartilage responses during early PTOA. A) Schematic of the knee joint illustrating the four histologic regions of interest (ROIs) used for Total Joint (dashed lines; B-D) and Medial Compartment (solid lines; G-I) semiquantitative OARSI scoring. B) Male Cx43^cKO^ (KO) mice had worse (higher score) total surface damage than Cx43^WT^ (WT) males (p < 0.0001), and KO females (p = 0.006), whereas females showed no genotype-dependent differences. C) Cartilage was thinner (total joint score) in KO males compared with WT (p = 0.007). D) Total hypertrophic chondrocyte (HC) scores. Female KO mice exhibited fewer HC than male KOs (p = 0.01). E-F) Representative SOFG-stained medial femur (top) and medial tibia (bottom) sections from E) male and F) female WT and KO mice. Insets (dashed boxes) highlight the articular surfaces of the medial compartment ROIs. KO males show pronounced articular surface damage and cartilage thinning within the medial joint compartment, whereas females demonstrate relatively preserved surface morphology and fewer HC in KO cartilage. G) KO males had worse surface damage scores than WT males at both medial sites (femur p < 0.0001; tibia p = 0.01), while females had no significant genotype differences. H) Male KO mice had thinner medial femoral and tibial cartilage compared with WT controls (p < 0.0001 and p = 0.01, respectively) without any genotype effect in females. I) Female KOs had fewer HC than WT females in both medial sites (femur p = 0.01; tibia p = 0.04), while no genotype effect was present in males. Additionally, in media tibia cartilage, female KOs had fewer HCs than female WT or male KOs (p = 0.03). Data is displayed as mean ± SD. *p<0.05, **p<0.01, ***p<0.001, ****p<0.0001.

To better understand the basis of sex-specific differences in OA severity observed in this model, we analyzed the femoral and tibial surfaces of injured medial joint compartment, as previously reported.^24,32,33^ We found that structural differences (i.e. surface damage and thickness) between KO and WT males were further highlighted; in male Cx43^cKO^ mice, both the medial femur (MF) and tibia (MT) had worse surface damage (genotype effect: F(1,32) = 21.4, p = 0.003, MF: Δ = −2.2, 95% CI: −3.08 to −1.32, p < 0.0001; MT: Δ = −1.1, 95% CI: −2.01 to −0.25, p = 0.01) and reduced cartilage thickness (genotype effect: F(1,32) = 19.2, p = 0.0001, MF: Δ = −1.7, −2.5 to −1, p < 0.0001; MT: Δ = −1.1, 95% CI: −1.9 to −0.4, p = 0.004) compared to WT (Figure 3G-I). Within Cx43^cKO^ males, the medial femur was more severely affected than medial tibia in both measures (surface damage Δ = 1.5, 95% CI: 0.4 to 2.7, p = 0.006; thickness Δ = 1.5, 95% CI: 0.45 to 2.5, p = No such genotype-associated differences were observed in the medial compartment of female mice with respect to surface damage (MF: p = 0.9, MT: p = 0.1) or cartilage thickness (MF: p = 0.2, MT: p = 0.6, Figure 3G-I). Further, KO males had worse surface damage scores for the medial femur compared to KO females (Δ = 1.3, 95% CI: 0.1 to 2.4, p = 0.03).

On the cellular level, female Cx43^cKO^ mice had a lower total hypertrophic chondrocyte (HC) score compared with KO males (Δ = 3.9, 95% CI: 0.93 to 6.87 p = 0.01), an effect most pronounced in the medial tibia (Δ = 1.0, 95% CI: 0.1 to 10.9, p = 0.03). Genotype also had an effect on chondrocyte responses in females; Cx43^KO^ females exhibited fewer HCs in both the medial femur and tibia compared to Cx43^WT^ females (MF: Δ = 0.9, 95% CI: 0.2 to 1.6, p = 0.01; MT: Δ = 0.7, 95% CI: 0.03 to 1.4, p = 0.04; Figure 3I). No genotype effect on HCs was observed in males. Data was analyzed by a two-way ANOVA and is displayed as mean differences and 95% CI, n = 5 per group.

### Cx43 knockout in cartilage induces sex-specific subchondral bone alterations in the destabilized region

To assess whether cartilage-specific Cx43 deletion alters periarticular and subchondral bone remodeling following joint injury, we performed quantitative micro–computed tomography (µCT) and complementary histological analyses at 12 weeks post-injury. Histological evaluation revealed that Cx43^KO^ females developed worse periarticular osteophyte (higher total scores) than Cx43^WT^ females (genotype effect (F1,16) = 5.8, p = 0.03, Δ = −2.5, 95% CI: −4.9 to −0.1, p = 0.04), with the medial tibia most severely affected (genotype effect (F1,16) = 10.4, p = 0.005, mean Δ = −1.00, 95% CI: −1.9 to −0.1, p = 0.02, Figure 4).

**Figure 4.**
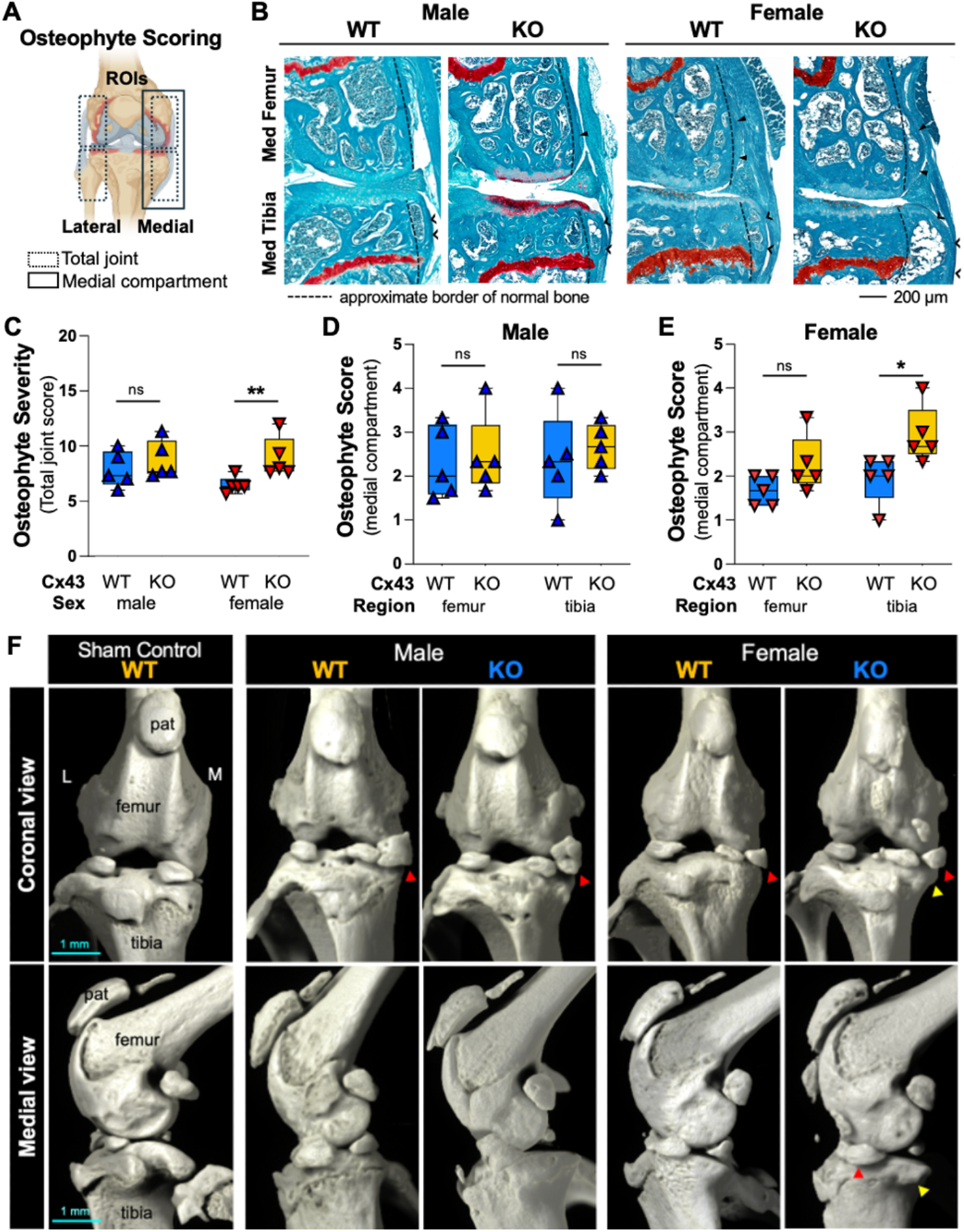
Cx43 knockout resulted in more severe osteophytes in females, but not males. A) Schematic of murine knee joint indicating four regions of interest (ROIs) scored on histology (sum expressed as whole-joint osteophyte score), and two regions comprising the medial compartment. B) Representative SOFG-stained histological images of osteophytes of the medial femoral condyle (open arrowheads) and tibial condyle (closed arrowheads) in male (left) and female (right) mice at 12-weeks post-DMM. C) Total osteophyte severity scores in DMM-operated limbs at 12 weeks post-injury, stratified by sex and genotype; Cx43^cKO^ (KO) females had larger osteophytes than Cx43^WT^ (WT) females (*p* = 0.004), with no genotype effect in males. D) The medial joint compartment in male mice (left), showed no difference in osteophytes severity between genotypes. However, in E) females, KOs had worse medial tibial osteophytes than WT females (*p* = 0.02). F) Representative 3D μCT coronal (top row) and medial (bottom row) view reconstructions of a sham-operated WT control (far left) and DMM-operated, male and female knee joints. All DMM joints exhibited extrusion of the anterior horn of the medial (M) meniscus with varying degrees of ossification (red markers) and periarticular osteophytes, especially notable in the medial tibial condyle of female KOs (yellow marker). Data (n=5/group) analyzed by two-way ANOVA, displayed as mean ± SD.*p<0.05, **p<0.01

Subchondral bone changes, analyzed by μCT, emerged later than cartilage alterations, becoming evident at 12 weeks post-DMM. Main genotype effect was significant in both the medial femur (F(1,8) = 12.55, p = 0.008) and medial tibia (F(1,8) = 7.50, p = 0.03). In females, Cx43^cKO^ mice showed reduced bone volume fraction (BV/TV) compared with Cx43^WT^ females in both the medial and lateral joint compartments (MF: Δ = −0.06, 95% CI: −0.1 to −0.02, p = 0.006, MT: Δ = −0.10, 95% CI: −0.2 to −0.01, p = 0.03). Further, BV/TV was lower in Cx43^cKO^ females compared with Cx43^cKO^ males (MF: Δ = −0.06, 95% CI: −0.1 to −0.02, p = 0.008, MT: Δ = −0.11, 95% CI: −0.2 to −0.03, p = 0.02, Figure 5).

**Figure 5.**
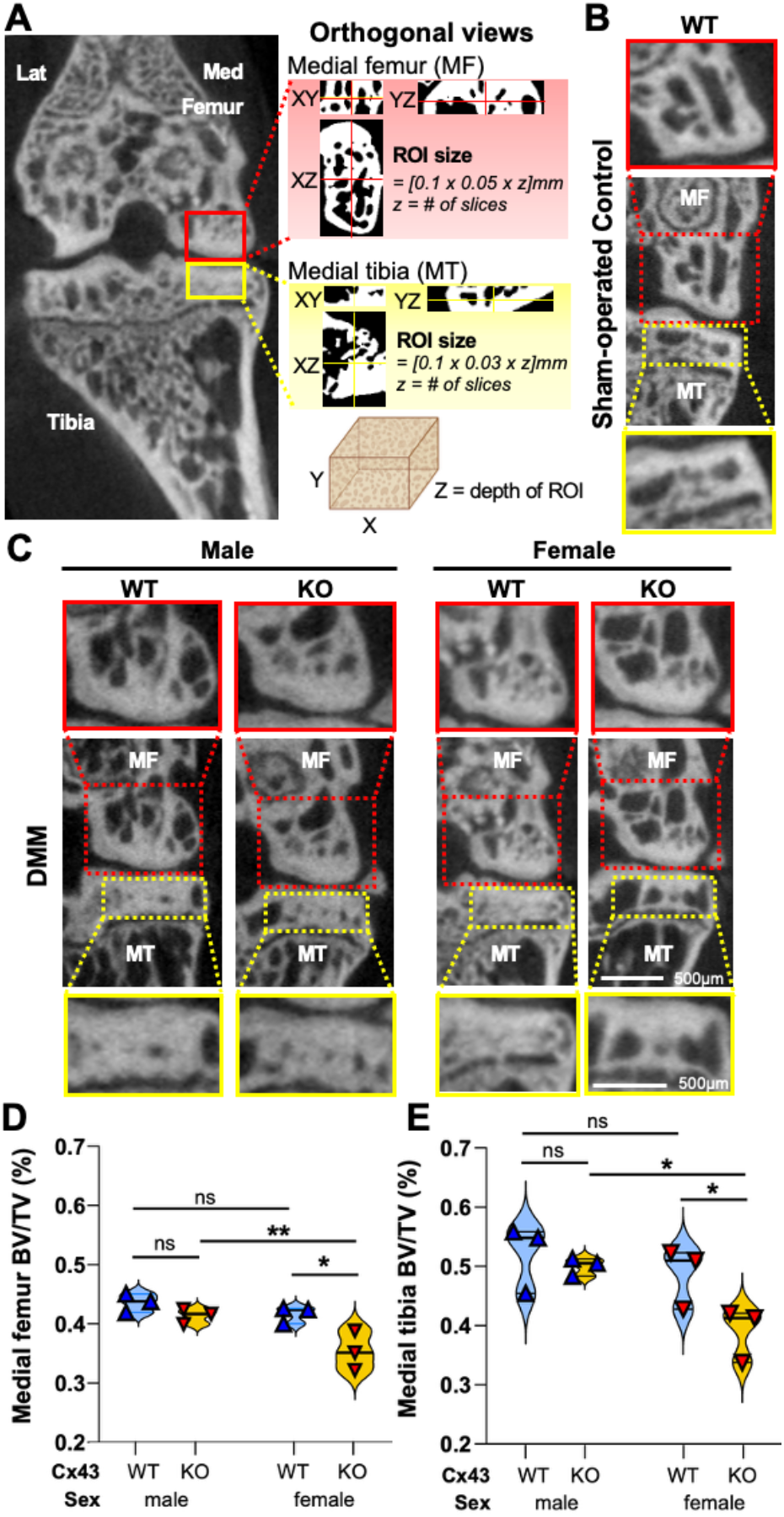
Cx43 cartilage knockout induces trabecular rarefaction in female DMM limbs. A) Schematic of μCT analysis showing regions of interest (ROIs) in the medial femur (MF) and medial tibia (MT), with representative coronal, sagittal, and transverse views. ROI size was calculated as 0.1 × 0.05 × z mm for MF and 0.1 × 0.03 × z mm for MT, where *z* = number of stack slices. B) Representative sagittal μCT images of subchondral bone from Cx43^WT^ (WT) female sham-operated limb, and C) WT and Cx43^cKO^ (KO) males and females 12 weeks post-DMM. Insets detail trabecular architecture differences between genotypes and sexes. (D) Quantitative analysis of bone volume fraction (BV/TV) in D) MF and E) MT. KO females showed significant trabecular rarefaction in MF (KO vs WT females, *p* = 0.006; KO females vs KO males, *p* = 0.008) and MT (KO vs WT females, *p* = 0.01; KO females vs KO males, *p* = 0.02), consistent with reduced load-bearing bone mass in female KO mice. Data (n=3/group) analyzed by two-way ANOVA, displayed as mean ± SD. *p<0.05, **p<0.01

### Cx43 Loss Impairs Bioenergetic Capacity and Mitochondrial Stress Responses in Human Chondrocytes

Given emerging evidence that Cx43 participates in mitochondrial regulation and stress adaptation in other tissues, we next sought to determine whether the structural and degenerative phenotypes observed in our cKO mice in vivo might be driven, in part, by a cell-intrinsic defect in chondrocyte bioenergetics. To test this directly and independent of joint-level and mechanical effects, we assessed mitochondrial function in immortalized human articular chondrocytes with CRISPR-mediated deletion of *GJA1* using Seahorse XF Mito Stress Tests (n = 9 per group, Figure 6A-F).

**Figure 6.**
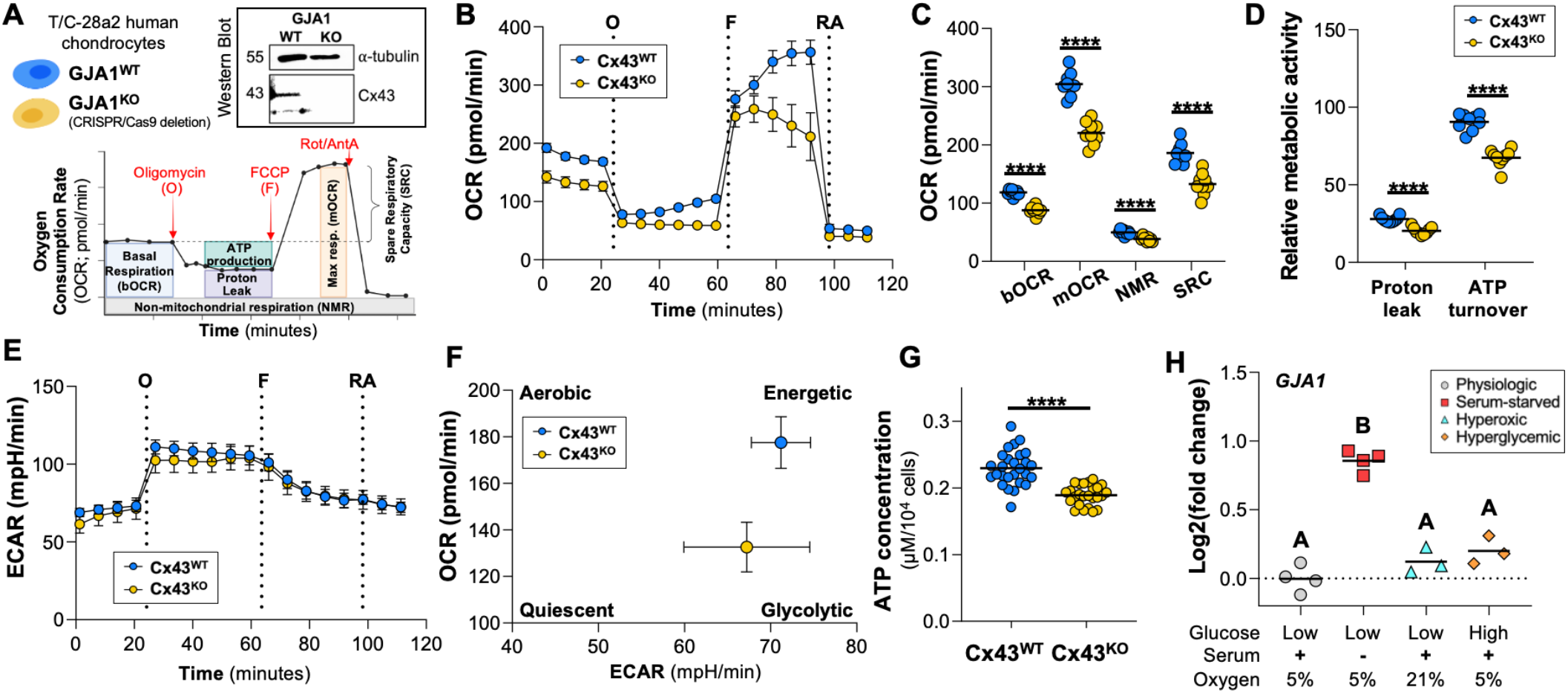
Loss of Cx43 impairs mitochondrial respiration and reduces ATP production in chondrocytes. A) Methods: Human Cx43 knockout (GJA1^KO^) chondrocyte line was validated via western blot and RT-qPCR. Mitochondrial function was assessed by microrespirometry, mitochondrial stress test and ATP luminescence assays. Schematic of mitochondrial stress test (bottom) using oligomycin (O), FCCP (F), and rotenone/antimycin A (RA) to determine chondrocyte stress responses. B) Oxygen consumption rate (OCR) profiles of Cx43^KO^ and Cx43^WT^ chondrocytes. Compared to WT chondrocytes, Cx43^cKO^ cells exhibited C) reduced basal respiration (bOCR), maximal respiration (mOCR), spare respiratory capacity (SRC), and D) proton leak and ATP turnover. E) Extracellular acidification rate (ECAR) profiles reflect no differences in glycolytic activity. F) Metabolic phenotype map (OCR/ECAR ratio) indicates that Cx43^KO^ chondrocytes are less aerobic and energetic than Cx43^WT^. H) Luminescence-based ATP assay confirmed lower ATP levels in Cx43^KO^ chondrocytes (n = 26/group). H) *GJA1* mRNA expression increased under serum starvation (0% serum) but was unchanged by hyperoxia (21% O_2_) or hyperglycemia (1 g/L glucose), suggesting regulation by glucose-independent metabolic stress. Data are presented as individual values with lines indicating group means. Statistical significance was assessed using t-tests or one-way ANOVA as appropriate, *p < 0.05, ****p < 0.0001.

Consistent with this hypothesis, Cx43-deficient chondrocytes (Cx43^KO^) exhibited a marked reduction in mitochondrial respiratory capacity across all measured parameters (Figure 6CD). Basal oxygen consumption rate (bOCR) was reduced by 26% in Cx43^KO^ cells compared to Cx43^WT^ controls (Δ = −30.7 pmol/min, 95% CI: −36.8 to −24.7; t(16) = 10.70, p < 0.0001, η^2^ = 0.88). Maximal oxygen consumption rate (mOCR) was decreased by 28% (Δ = −83.9 pmol/min, 95% CI: −104.0 to −63.9; t(16) = 8.86, p < 0.0001, η^2^ = 0.83) indicating a reduced capacity to respond to energetic demand. Consistent with this, spare respiratory capacity (SRC) was reduced by 29% in Cx43^cKO^ chondrocytes (Δ = −53.2 pmol/min, 95% CI: −71.0 to −35.4; t(16) = 6.35, p < 0.0001, η^2^ = 0.72), suggesting limited metabolic flexibility under stress conditions. ATP-linked respiration was also decreased by 26% (Δ = −23.1 pmol/min, 95% CI: −28.6 to −17.7; t(16) = 8.98, p < 0.0001, η^2^ = 0.83), consistent with impaired oxidative phosphorylation and decreased cellular energy production.

Non-mitochondrial respiration (NMR) was reduced in Cx43^KO^ chondrocytes (Δ = −11.6 pmol/min, 95% CI −15.7 to −7.4; t(16) = 5.90, p < 0.0001), and proton leak was decreased by 27% in Cx43^KO^ cells relative to WT (Δ = −7.63 pmol/min, 95% CI −9.87 to −5.39; t(16) = 7.23, p < 0.0001), consistent with reduced basal mitochondrial activity rather than increased uncoupling. Extracellular acidification rate (ECAR) profiles revealed no significant differences in glycolytic activity between WT and Cx43^KO^ cells (Figure 6E). However, metabolic phenotype mapping demonstrated a shift toward a less energetic, more quiescent metabolic state in Cx43-deficient chondrocytes (Figure 6F), characterized by reduced overall bioenergetic output without compensatory upregulation of glycolysis.

Consistent with this global reduction in metabolic activity, total cellular ATP content was significantly decreased in Cx43^KO^ cells as measured by luminescence assay (n = 25–27 per group; mean difference = 0.04 ± 0.01 nmol/10^4^ cells; p < 0.0001, Figure 6G). Together, these data demonstrate that loss of Cx43 is associated with a coordinated reduction in cellular bioenergetic capacity in human chondrocytes, supporting a cell-intrinsic role for Cx43 in maintaining metabolic homeostasis.

To explore whether *GJA1* expression is transcriptionally regulated under metabolic stress, mRNA levels were assessed in immortalized human chondrocytes exposed to serum starvation, hyperoxia, or high glucose (Figure 6H n = 3-4). Treatment had a significant overall effect on *GJA1* expression (one-way ANOVA, F(3,10) = 68.61, p < 0.0001), which was upregulated under serum starvation conditions (mean difference = −0.86, 95% CI: −1.1 to −0.7, p < 0.0001) whereas no differences were detected between physiologic and hyperoxic or hyperglycemic conditions (all p > 0.05; Figure 6H). These findings indicate *GJA1* expression is responsive to nutrient deprivation.

## Discussion

The present study demonstrates that cartilage-specific deletion of connexin 43 (Cx43; *Gja1*) in mice accelerates PTOA progression in a sex-dependent manner and impairs mitochondrial bioenergetic capacity in human chondrocytes. Together, these findings identify Cx43 as a previously underappreciated regulator of cartilage metabolic resilience following injury. Although increased Cx43 expression has been associated with inflammatory and senescence-associated phenotypes in OA cartilage,^20,34^ our data indicate that complete loss of Cx43 within cartilage compromises structural homeostasis and exacerbates joint degeneration after mechanical injury. These findings highlight an important distinction between partial modulation of Cx43 signaling and complete genetic ablation, with direct implications for the development of connexin-targeted therapies.

Cx43 is a central mediator of intercellular communication in musculoskeletal tissues and contributes to mechanotransduction, extracellular matrix (ECM) regulation, and coordinated cellular stress responses.^14,18,35–37^ In cartilage, prior in vitro studies demonstrated that disruption of Cx43 signaling alters matrix synthesis, gap-junction communication, and chondrocyte differentiation.^38–41^ Similarly, truncation of the C-terminal regulatory domain of Cx43 results in abnormal cartilage architecture and accelerated degeneration.^38^ Collectively, these observations support a model in which tightly regulated Cx43 activity is required for maintenance of cartilage homeostasis, whereas complete loss of Cx43 impairs the ability of cartilage to adapt to mechanical and metabolic stress.

A notable finding of the present study was the sexually dimorphic response to cartilage-specific Cx43 deletion. Female Cx43^cKO^ mice exhibited earlier global joint degeneration, reduced hypertrophic chondrocyte abundance, increased osteophyte formation, and subchondral trabecular rarefaction, whereas male knockouts developed more severe cartilage surface disruption and thinning. These findings suggest that loss of Cx43 alters osteochondral adaptation to injury differently in male and female joints. Although the mechanisms underlying these differences remain incompletely defined, prior studies indicate that sex hormones modulate both chondrocyte maturation and Cx43-mediated signaling.^42,43^ Estrogen enhances Cx43-dependent mechanosensitivity and gap-junction communication in osteocytes,^42^ while osteocyte-specific deletion of Cx43 alters bone remodeling dynamics and increases cortical bone resorption. Our findings extend these observations to the osteochondral unit and suggest that sex-specific regulation of Cx43 signaling may influence how cartilage and bone respond to mechanical injury.

Importantly, despite cartilage-restricted deletion of Cx43, female cKO mice exhibited substantial changes in periarticular and subchondral bone remodeling, including increased osteophyte formation and reduced subchondral bone volume fraction. These findings support the concept that cartilage-derived signals strongly influence post-injury bone adaptation. Altered cartilage mechanics, extracellular matrix integrity, inflammatory mediator release, or osteochondral biochemical signaling could all contribute to the divergent bone remodeling phenotype observed in Cx43-deficient joints. Previous studies have established Cx43 as a key regulator of skeletal cell communication and mechanobiology within bone.^18,35,44^ The present findings suggest that disruption of Cx43 signaling within cartilage alone is sufficient to perturb osteochondral crosstalk during PTOA progression.

Our bioenergetic studies further identify Cx43 as an important regulator of chondrocyte mitochondrial function. Cx43-deficient human chondrocytes exhibited coordinated reductions in basal respiration, maximal respiration, spare respiratory capacity, ATP-linked respiration, and total ATP production, without compensatory increases in glycolysis. These data indicate that loss of Cx43 drives a metabolically constrained or quiescent phenotype characterized by impaired oxidative phosphorylation and reduced energetic flexibility. Because chondrocytes experience fluctuating mechanical, inflammatory, and nutritional stress within the injured joint, the inability to mount an effective mitochondrial stress response may contribute directly to the accelerated cartilage degeneration observed in vivo.

These findings align with emerging evidence that Cx43 and the alternatively translated GJA1-20k isoform regulate mitochondrial dynamics and stress adaptation in non-musculoskeletal tissues.^21,22,45,46^ In cardiomyocytes, mitochondrial-localized Cx43 supports mitochondrial network organization, preserves complex I activity, and promotes metabolic adaptation during ischemic stress.^21,22,45^ Although the precise mechanisms linking Cx43 to mitochondrial regulation in chondrocytes remain unclear, our findings extend these observations to articular cartilage and support a model in which Cx43 contributes directly to maintenance of cellular metabolic competence. Notably, this metabolic role may help reconcile apparently conflicting observations in the OA literature. While elevated Cx43 expression has been linked to inflammatory and senescence-associated phenotypes in diseased cartilage,^19,20^ mitochondrial Cx43 signaling has also been shown to support stress adaptation and cell survival in other tissues.^21,22,45^ Thus, the biologic effects of Cx43 may depend strongly on cellular context, subcellular localization, and the degree of signaling disruption.

Consistent with this concept, we observed that GJA1 expression increased during serum starvation but not under hyperoxic or hyperglycemic conditions, suggesting that nutrient stress may selectively regulate Cx43 expression in chondrocytes. Previous studies similarly demonstrated that inflammatory cytokines and growth factors modulate Cx43 turnover and gap-junction signaling in cartilage.^14,41,47^ Together, these findings support the hypothesis that Cx43 functions as part of an integrated cellular stress-response network linking metabolic state, mechanotransduction, and intercellular communication within the joint.

Several limitations should be acknowledged. First, only a single PTOA model and two post-injury timepoints were evaluated, limiting assessment of the full temporal evolution of Cx43-dependent joint remodeling. Second, although the Agc1-CreERT2 system enabled targeted deletion in aggrecan-expressing cartilage tissues, this model does not isolate potential indirect effects arising from meniscal or other cartilaginous tissues. Third, mitochondrial studies were performed in an immortalized human chondrocyte line, which differs metabolically from primary articular chondrocytes despite retaining functional gap-junction signaling and relevant receptor expression.^39,48^ Finally, although our findings strongly implicate altered osteochondral communication, direct effects on bone cell populations cannot be resolved using the present cartilage-specific model and will require complementary approaches.

## Conclusion

This study demonstrates that cartilage-specific loss of Cx43 accelerates posttraumatic OA in a sex-dependent manner and impairs mitochondrial bioenergetics in human chondrocytes. These findings identify Cx43 as a key regulator of metabolic and structural adaptation and suggest that Cx43-dependent stress responses may be important for maintaining cartilage resilience after joint injury. From a therapeutic perspective, our data argue against global suppression of Cx43 activity in OA and instead support strategies that selectively modulate connexin signaling or stabilize Cx43-associated mitochondrial functions. Finally, this work suggests a more granular understanding of Cx43 biology in the context of joint disease may lead to new therapeutic strategies to prevent and treat OA.

## Materials and Methods

### Generation of the Cx43cKO Mouse Line

Cartilage-specific Cx43 knockout mice were generated by crossing cartilage-specific Cre transgenic mice (B6.Cg-Acan^tm1(cre/ERT2)Crm/J^, #019148) with homozygous floxed Gja1 mice (B6.129S7-Gja1^tm1Dlg^/J, #008039; both The Jackson Laboratory). F1 offspring (Gja1^flox/+^; Cre^+/−^) were backcrossed to the floxed Gja1 line to obtain F2 mice homozygous for the floxed allele and either Cre-positive (Cx43^cKO^) or Cre-negative (Cx43^WT^) genotypes (Figure 1A). Cre-positive mice enable tamoxifen-inducible deletion of Gja1 in aggrecan-expressing cartilage tissues. The colony was maintained by breeding Gja1^flox/flox^; Cre^+^ mice with Gja1^flox/flox^; Cre^-^ littermates. Genotyping was performed on tail snips collected at weaning (Transnetyx, Inc.). All animals were housed under standard conditions in the institution’s Center for Animal Research and Education (CARE) facility, and all breeding and surgical procedures were approved by the institution’s IACUC (protocol #2019-0045, 2019-0010). For breeding details, see Supplementary Methods.

### Induction of conditional Cx43 knock-out

To induce Cre-mediated *Gja1* recombination, 3-month-old Cre+ and Cre-control mice received tamoxifen (TMX, 75 mg/kg IP) once daily for 5 days (Figure 1B). TMX was prepared as a 30 mg/ml TMX-free base (Krackeler Scientific, 45-T5648-5G-EA) solution in ethanol/sunflower oil (1:9), aliquoted, and stored at −20 °C. Injections (1ml low dead-space syringe, 27G ½” needle) were alternated between left and right lower abdominal quadrants. Mice were housed in clean cages beginning 7-days post-injection to minimize TMX exposure during subsequent procedures.

### Validation of conditional Cx43 knock-out

Conditional deletion of Gja1 was confirmed at the transcript and protein levels using RT-qPCR, immunohistochemistry, and Western blotting. For transcript analysis, mRNA was isolated from articular cartilage, heart, and kidney of tamoxifen-induced Cx43^cKO^ and Cx43^WT^ mice and Gja1 expression was quantified by RT-qPCR using GAPDH as the reference gene. Cx43 protein loss within cartilage was assessed by immunohistochemistry on knee joint sections collected 7 days after tamoxifen induction, using a C-terminal anti-Cx43 antibody and DAB detection. DAB-positive area was quantified in ImageJ. Tissue-specific deletion was further evaluated by Western blotting of Cx43 in articular cartilage, xiphoid cartilage, heart, and kidney from adult Cx43^cKO^ and Cx43WT mice. Together, these assays verified efficient and cartilage-specific deletion of Gja1 following tamoxifen administration. All assays are detailed in Supplementary Methods.

### PTOA model and histological assessment

Post-traumatic osteoarthritis (PTOA) was induced in 3-month-old Cx43^cKO^ and Cx43^WT^ mice using the well-established DMM model^49^ (Figure 2B). Mice were divided into experimental groups based on genotype (Cx43^cKO^ or Cx43^WT^), sex (male or female), surgical procedure (DMM or sham), and postoperative interval (4 or 12 weeks), with 5 animals per subgroup (n = 5/group). The left hindlimb underwent DMM surgery, while the contralateral limb received a sham operation (arthrotomy without ligament transection, Figure 2B). Buprenorphine SR (Ethiqa XR, 3.25 mg/kg SC) was administered pre-operatively, with additional post-operative doses given based on individual clinical assessment. All DMM surgeries were performed by the same person. Mice were monitored daily for 5 days post-op and every 3-5 days thereafter. At 4- or 12-weeks post-surgery, animals were euthanized with CO_2_. Knee joints were dissected, fixed in 4% paraformaldehyde with 1% cetylpyridinium chloride for 24 h at 4 °C, and decalcified in 10% EDTA (pH 7.2–7.4) for 8 days.

Joints were paraffin-embedded, sectioned in the frontal plane at 5 μm thickness, and stained with Safranin-O/Fast Green. Images were acquired using a digital scanning microscope Aperio Scanscope. Articular cartilage degeneration was assessed in 4 regions of interest (ROI): medial femoral condyle (MF), medial tibial condyle (MT), lateral femoral condyle (LF) and lateral tibial condyle (LT). An established semi-quantitative OARSI scoring rubric^27^ was used to generate one total joint score and subsequently modified to include cellular, tissue integrity and subchondral bone parameters (Table 1). Histological grading was performed by two blinded DVM observers, and results were expressed as mean ± SD. The expression of Cx43 at 4- and 12-weeks post-injury was investigated via IHC using the same staining protocol as for the *Gja1* knockout validation. An overview of study timeline and experimental groups is illustrated in Figure 2.

### Radiographic analysis

At study endpoints (4- and 12-weeks), hind limbs (n = 3 per group) were collected and scanned using a Bruker SkyScan 1276 micro-CT system at 10 µm voxel resolution with 70 kV, 200 µA, and typical exposure time of 770ms. Image reconstruction was performed in Dragonfly software (Object Research Systems, Montreal, Canada). Bone volume fraction (BV/TV) was quantified in the medial femoral condyles (MF) and tibial plateaus (MT) using ImageJ (v1.53) and BoneJ plugin. Regions of interest were manually selected using consistent anatomical landmarks in both MF and MT (Figure 4). Detailed methodology is displayed in Supplementary Materials.

### Chondrocyte bioenergetics

To investigate the role of Cx43 in mitochondrial metabolism, Cx43^KO^ human chondrocytes were generated from an immortalized human chondrocyte line (T/C-28a2; ATCC® CRL-2846™) using CRISPR/Cas9 genome editing as previously described.^20^ Knockout efficiency was validated by Western blotting. Cx43^WT^ and Cx43^KO^ chondrocytes were cultured under four distinct metabolic conditions for 24 hours: (1) baseline [0.45 g/L glucose, 10% fetal bovine serum (FBS), 5% O_2_], (2) serum starvation [0% FBS], (3) hyperoxia [21% O_2_], and (4) hyperglycemia [1.0 g/L glucose]. Total RNA was isolated and analyzed by RT-qPCR to assess transcriptional responses.

Mitochondrial respiration was assessed using high throughput microrespirometry (XFe96 Analyzer, Seahorse Biosciences, Figure 6B). Cx43^WT^ and Cx43^cKO^ chondrocytes were plated in Seahorse XF96 cell culture microplates and subjected to a mitochondrial stress test, measuring oxygen consumption rate (OCR) under basal conditions and in response to oligomycin, FCCP, and a combination of rotenone and antimycin A. Bioenergetic parameters including basal respiration, ATP production, proton leak, and spare respiratory capacity were calculated according to manufacturer’s guidelines (Agilent Seahorse XF Cell Mito Stress Test). Each condition was tested in 6-8 biological replicates across three independent experiments.

Intracellular ATP levels were quantified using a luminescent ATP detection kit (Abcam ab113849). Substrate and ATP standards were reconstituted per manufacturer instructions, and an 8-point standard curve (0–10 µM) was prepared in culture media. Cells were adjusted to 100 µL/well, lysed with 50 µL detergent, and incubated with 50 µL substrate solution. Plates were shaken for 5 min after each reagent addition, dark-adapted for 10 min, and luminescence was measured without attenuation. ATP concentrations were calculated from the standard curve and normalized to cell number or protein content as indicated.

### Statistical methods

A priori power analysis was performed based on published OARSI histology data from the mouse DMM model^27^ and n = 5 limbs per group (total 40 mice; 80 limbs) were used (Figure 2C). Details of the power analysis are provided in the Supplementary Materials. All statistical analyses were performed using JMP Pro (v17.0.0, SAS Institute) and GraphPad Prism (v10.3.1).

For qPCR validation of *Gja1* deletion, a one-tailed t-test was used based on an *a priori* hypothesis of reduced transcript abundance in Cx43^cKO^ tissue, whereas IHC data were analyzed using two-tailed nonparametric tests due to non-normal data distributions. Quantitative IHC data mapping Cx43 expression in sham- and DMM-operated limbs over time were analyzed using linear mixed-effects models with surgical condition and timepoint as fixed effects and animal ID as a random effect. Type III F tests were used, followed by Fisher’s least significant difference post hoc tests where appropriate.

OARSI whole-joint scores and modified OARSI feature-level scores were analyzed using two-way analysis of variance (ANOVA). Fixed factors included surgery (sham vs DMM), genotype (Cx43^WT^ vs Cx43^cKO^), and sex, with sex included either as an interaction term or by stratified analysis as indicated in the Results. Micro–computed tomography (μCT) outcomes, including bone volume fraction (BV/TV), were analyzed using two-way ANOVA with genotype and sex as fixed factors. All tests were two-tailed, and exact statistics, degrees of freedom, and sample sizes are reported in the Results and corresponding figure legends.

For bioenergetic analyses, two tailed unpaired t-tests were used. For transcriptional responses to metabolic stress, one-way ANOVA followed by Tukey’s multiple comparisons test was used. P < 0.05 was considered statistically significant.

Figures were created using Microsoft PowerPoint (v 2511) and BioRender.com.

## ACKNOWLEDGEMENTS

We thank Dr. Purva Singh and Dr. Camila Carballo for training and guidance on histological scoring, Matthew Thomas for technical support, Teresa Porri for assistance with micro-CT image acquisition, Cornell’s Center for Animal Resources and Education for support with mouse colony management, and the Cornell Statistical Consulting Unit for statistical expertise.

## Ethics/Compliance

All experiments involving human chondrocyte cell lines and gene editing were conducted under approval of the Institutional Biosafety Committee (IBC; MUA #16342-2), with work performed at BSL-2 in compliance with NIH and institutional guidelines. The human immortalized cell line (T/C-28a2; ATCC® CRL-2846™) was provided by Dr. Miguel Otero from the Hospital for Special Surgery, New York, NY. Human T/C-28a2 Cx43^KO^ chondrocytes were provided by Dr. Maria Mayan, CellCOM Research Group, Center for Research in Nanomaterials and Biomedicine (CINBIO), University of Vigo, Spain.

## Data availability

The datasets generated during and/or analyzed during the current study are available from the corresponding author on reasonable request.

## Author contributions

**Conceptualization:** Author 9. **Resources:** Author 3, 8, 9. **Data curation:** Author 1, 2, 3, 4, 5, 6, 7, 9. **Formal analysis:** Author 1, 2, 3, 9. **Funding acquisition:** Author 9. **Investigation:** Author 1, 2, 3, 4, 6. **Methodology:** Author 1, 2, 3, 8, 9. **Project administration:** Author 9. **Software:** Author 1, 3. **Supervision:** Author 9. **Validation:** Author 1, 2, 4 **Visualization:** Author 1, 2, 3, 4, 5, 6, 9. **Writing – original draft:** Author 1, 2. **Writing – review & editing:** All authors.

