## Supplemental Material for "Loss of connexin 43 in cartilage causes mitochondrial dysfunction and accelerates post-traumatic osteoarthritis progression"

### SUPPLEMENTARY MATERIALS

#### 1. Supplementary Methods

##### Generation of the Cartilage-Specific Cx43 Conditional Knockout Mouse Line

##### Breeding Strategy

Cartilage-specific Cx43 conditional knockout (Cx43cKO) mice were generated using an Agc1-Cre$^{ERT2}$ driver to induce tamoxifen-dependent recombination in aggrecan-expressing cells. Agc1-Cre$^{ERT2}$ transgenic mice (B6.Cg-Acan$^{tm1(cre/ERT2)Crm/J}$, The Jackson Laboratory, stock #019148) were crossed with homozygous floxed Gja1 mice (B6.129S7-Gja1$^{tm1Dlg}$/J, stock #008039) to produce F1 progeny (Gja1^flox/+; Cre^+/−). F1 offspring were then backcrossed to the Gja1^flox/flox line to generate F2 mice homozygous for the floxed Gja1 allele and either Cre-positive (Gja1$^{flox/flox}$; Cre$^+$) or Cre-negative (Gja1$^{flox/flox}$; Cre$^-$) genotypes. Cre-positive F2 mice were used as experimental Cx43$^{cKO}$ animals, while Cre-negative littermates served as controls (Cx43$^{WT}$). The colony was maintained by breeding Gja1$^{flox/flox}$; Cre$^+$ mice with either Gja1$^{flox/flox}$; Cre$^-$ or Gja1$^{flox/flox}$ mice to ensure consistent generation of both genotypes.

##### Genotyping

At 14-20 days of age, all pups were identified via paw tattoo and a distal tail biopsy was collected for genomic analysis. DNA extraction and automated genotyping were performed by Transnetyx, Inc. using custom-designed probes for the floxed Gja1 allele and the Agc1-CreERT2 transgene.

##### Animal Housing and Husbandry

All mice were housed in microisolator cages within the Cornell University Center for Animal Resources and Education (CARE). Environmental conditions were maintained at 21.1–22.7 °C, 35–45% humidity, and a 14 h light/10 h dark photoperiod. Mice had ad libitum access to standard rodent chow and water. Health monitoring followed CARE sentinel and pathogen surveillance protocols.

##### Immunohistochemistry validation of Cx43 cKO

To confirm *Gja1* deletion efficiency, immunohistochemistry for Cx43 was performed on knee joints at 7 days post-tamoxifen administration (n = 3-8 per genotype, 3 months of age). Formalin-fixed, paraffin-embedded sagittal sections were processed using sequential antigen retrieval with proteinase K (25530015, ThermoFisher Sicentific, 20ug/ml, 10min, RT) and hyaluronidase (H3506, Sigma Aldrich, 1600 U/mL, 20min, 37°C), followed, followed by quenching, blocking, and overnight incubation with a C-terminal polyclonal rabbit anti-Cx43 antibody (1:50 dilution; Thermo Fisher, 710700). HRP-conjugated secondary antibody (goat anti-rabbit IgG HRP-conjugated, 1:1000; Fisher Scientific, PI31460) and DAB substrate were used for detection, with hematoxylin counterstaining. Whole-slide images were scanned (Aperio ScanScope) and analyzed in ImageJ. DAB-positive area was quantified in articular cartilage and meniscal regions using Color Deconvolution2 plugin and thresholding (RenyiEntropy algorithm). For each section, the cartilage or meniscal ROI was manually outlined based on tissue boundaries, and the total DAB-positive area within this ROI was measured. Surface area percentage was calculated as (DAB-positive area / total manually-selected tissue area) × 100). Cx43 signal was compared between genotypes and timepoints.

#### RT-qPCR validation of *Gja1* deletion

To evaluate *Gja1* expression at the transcript level, mRNA was isolated from articular cartilage, heart, and kidney tissues of TMX-injected $Gja1^{flox/flox}$; $Cre^{+}$ ($Cx43^{cKO}$) and $Gja1^{flox/flox}$; $Cre^{-}$ ($Cx43^{WT}$) mice (n = 3 months of age, Figure 1C, G). Tissues were homogenized in TRIzol (Invitrogen, 15596026) using a Precellys Evolution homogenizer with CK28 lysing kit (Bertin, P000911-LYSK1-A.0) RNA was purified (RNeasy Mini Kit, QIAGEN, 74106), quantified (NanoDrop, Thermo Fisher Scientific) and reverse-transcribed (MultiScribe™ RT, Thermo Fisher, 4311235) using random primers. Quantitative PCR was performed with PowerUp SYBR Green Master Mix (Thermo Fisher, A25742) on a QuantStudio 5 system (Applied Biosystems). Relative *Gja1* expression was calculated using the ΔΔCT method, normalized to GAPDH. All samples were run in technical triplicate.

#### Western blot Cx43 cKO validation

Western blotting was performed on articular cartilage (femoral head), xiphoid cartilage, heart, and kidney from $Cx43^{WT}$ and $Cx43^{cKO}$ 6 months old mice to assess *Gja1* protein expression and confirm tissue specificity of the knockout. Tissues were lysed in RIPA buffer with protease and phosphatase inhibitors, and total protein was denatured in LDS sample buffer with DTT. Proteins were separated by SDS-PAGE (4-12% Bis-Tris gels) and transferred to PVDF membranes using the iBlot2 system. Membranes were blocked and incubated with anti-Cx43 (1:2500, Sigma C6219) and anti-α-tubulin (1:2500, Abcam ab7291) primary antibodies, followed by fluorescent secondary antibodies. Blots were imaged on a Bio-Rad ChemiDoc MP system. Quantification was not performed, as the purpose was validation of knockout efficiency and specificity.

#### A priori power analysis

$\sigma = 1.5$, $\mu_e = 4.4$, $\mu_o = 0.5$, and a 3:1 allocation ratio. Using $Z_a = 3.29$ ($\alpha = 0.001$) and $Z_\beta = 1.64$ (power = 95%), a sample size of n = 5 limbs per group was determined to provide >90% power.

### 2. Supplementary Data

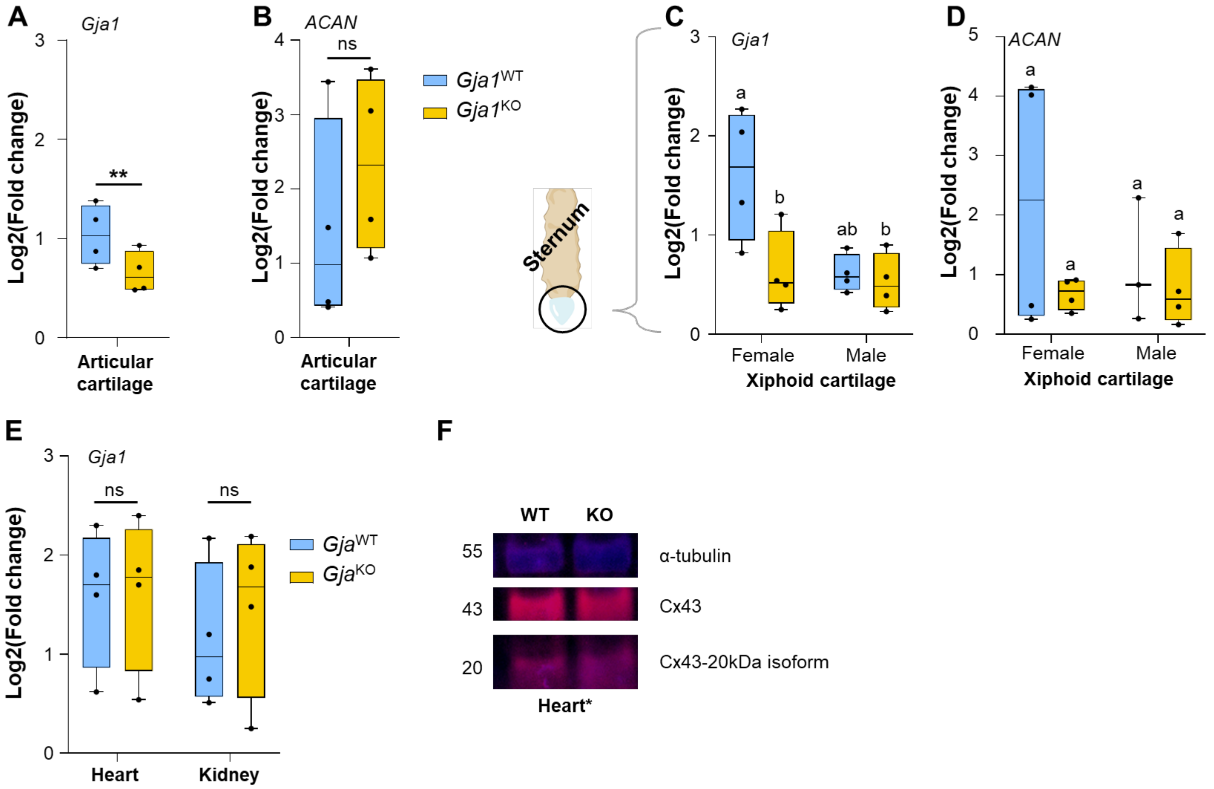


**Figure S1. Validation of tissue-specific Cx43 knockout in articular and xiphoid cartilage, heart and kidney at 12-weeks post-TMX injection**. A) RT-qPCR validation of tissue-specific recombination at 6 months of age confirms significant downregulation of Connexin43 gene (*Gja1*) expression in articular cartilage of Cx43$^{cKO}$ (*Gja1$^{KO}$;* yellow data) mice compared to Cx43$^{WT}$ (*Gja1$^{WT}$;* blue data). B) Aggrecan gene (*Acan*) expression in articular cartilage was not different between genotypes. C) In the xyphoid cartilage of 6 month old mice, Cx43$^{WT}$ females exhibited higher *Gja1* expression compared to Cx43$^{cKO}$, whereas Cx43$^{WT}$ and Cx43$^{cKO}$ males showed no difference ($p = 0.04$ for genotype effect). D) *Acan* expression in xiphoid cartilage was not different between genotype or sex. E) No genotype-dependent differences were observed in *Gja1* expression in non-cartilaginous tissues (heart and kidney). F) Western blot showing Cx43 and α-tubulin expression in heart tissue of Cx43$^{WT}$ and Cx43$^{KO}$ mice. Data shown as individual values with mean ± SD; Mann-Whitney test. *p < 0.05, **p < 0.01, ***p < 0.001, ****p < 0.0001.

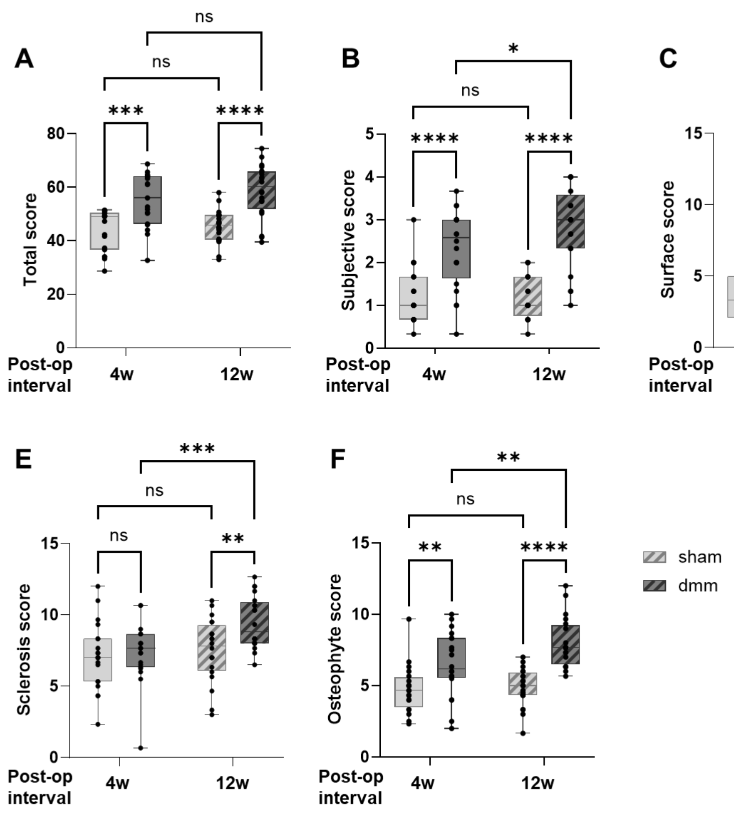


**Figure S2. Temporal changes in joint outcome measures following destabilization of the medial meniscus (DMM).** Box-and-whisker plots show A) Total and, B-F) composite histologic scores in sham-operated and DMM-operated knees at 4 weeks and 12 weeks post-operatively. Each panel represents a distinct joint parameter, with paired comparisons between sham and DMM limbs at each time point. Individual data points represent single knees; boxes indicate the interquartile range with median, and whiskers denote the full data range. Across multiple outcome measures, DMM-operated joints demonstrate progressive degenerative changes (increasing scores), consistent with the development and progression of post-traumatic osteoarthritic changes over time, whereas sham-operated joints do not.

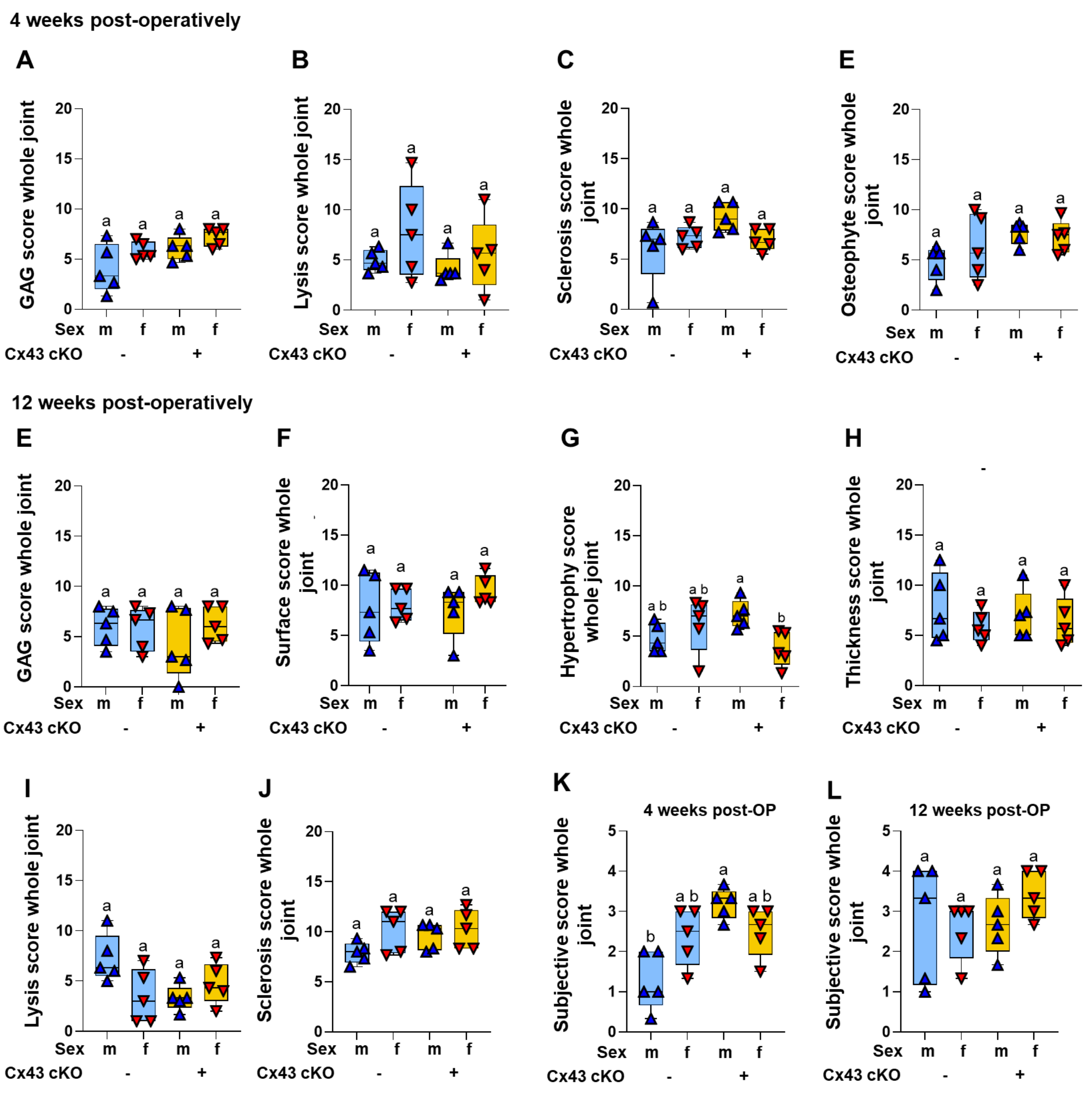


**Figure S3. Whole-joint histological outcomes at 4 and 12 weeks by Sex and Genotype.** Whole-joint histological scores are shown for male and female $Cx43^{WT}$ and $Cx43^{cKO}$ mice at 4 and 12 weeks post-DMM. A–D) Whole-joint component scores at 4 weeks; A) Whole-joint glycosaminoglycan (GAG) content, B) bone lysis, C) subchondral bone sclerosis, and D) osteophyte scores. E–H) Whole-joint outcome measures at 12 weeks; E) GAG content, F) surface quality, G) hypertrophic chondrocyte, H) cartilage thickness, I) subchondral bone lysis, and J) subchondral bone sclerosis. K-L) Subjective whole joint scores (0-4) at K) 4 weeks and L) 12-weeks post-DMM. Group differences were assessed by Kruskal-Wallis test with multiple comparisons; groups not sharing a letter are significantly different at $p<0.05$.
